# An Aging-Susceptible Circadian Rhythm Controls Cutaneous Antiviral Immunity

**DOI:** 10.1101/2023.04.14.536934

**Authors:** Stephen Kirchner, Vivian Lei, Paul Kim, Meera Patel, Jessica Shannon, David Corcoran, Dalton Hughes, Diana Waters, Kafui Dzirasa, Detlev Erdmann, Jörn Coers, Amanda MacLeod, Jennifer Y. Zhang

**Affiliations:** Department of Dermatology, Duke University, Durham, NC, USA; Department of Molecular Genetics and Microbiology, Duke University, Durham, NC, USA; Department of Immunology, Duke University, Durham, NC, USA; Duke Center for Genomic and Computational Biology, Duke University, Durham, NC, USA; Department of Neurobiology, Duke University, Durham, NC, USA; Department of Psychiatry and Behavioral Sciences, Duke University, Durham, NC, USA; Department of Biomedical Engineering, Duke University, Durham, NC, USA; Department of Neurosurgery, Duke University, Durham, NC, USA; Howard Hughes Medical Institute, Chevy Chase, Maryland 20815, USA; Department of Surgery, Duke University, Durham, NC, USA; Janssen Pharmaceuticals, San Diego, CA, USA; Department of Pathology, Duke University, Durham, NC, USA

**Author notes:** Corresponding author: Jennifer Zhang.

## Abstract

Aged skin is prone to viral infections, but the mechanisms responsible for this immunosenescent immune risk are unclear. We observed that aged murine and human skin expressed reduced antiviral proteins (AVPs) and circadian regulators including Bmal1 and Clock. Bmal1 and Clock were found to control rhythmic AVP expression in skin and such circadian-control of AVPs was diminished by disruption of immune cell interleukin 27 signaling and deletion of Bmal1/Clock genes in mouse skins, as well as siRNA-mediated knockdown of CLOCK in human primary keratinocytes. We found that treatment of circadian enhancing agents, nobiletin and SR8278, reduced infection of herpes simplex virus 1 (HSV1) in epidermal explants and human keratinocytes in a Bmal1/Clock-dependent manner. Circadian enhancing treatment also reversed susceptibility of aging murine skin and human primary keratinocytes to viral infection. These findings reveal an evolutionarily conserved and age-sensitive circadian regulation of cutaneous antiviral immunity, underscoring circadian restoration as an antiviral strategy in aging populations.

## INTRODUCTION

The skin acts as a physical barrier to invading pathogens, which can be disrupted by genetic defects, environmental challenges, wounds, and micro-injuries^1^. Skin barrier disruptions are of special concern for elderly patients due to the reduced regenerative capacity in aged skin. Consequently, these patients experience an increased risk of pathogen infection and other clinical issues^2, 3^. Nevertheless, how an aged skin microenvironment affects barrier function and immunosenescence is not well-understood. Aspects of the skin microenvironment that influence barrier defense include location of disruption, microbial content, moisture status, and age of the skin^3^. In addition, the time at which a wound is inflicted changes barrier responses and results in differential healing rates^4^, suggesting that the circadian rhythm in skin regulates tissue regeneration and immune responses.

The circadian rhythm controls time-of-day biological responses and regulates components of cell proliferation and wound re-epithelialization^5^. Mice deficient in Bmal1, one of the core transcription factors of the circadian clock, exhibit greater burden in viral infections ^6, 7^, indicating that circadian rhythms influence antiviral functions. Circadian function declines in older individuals^8^; however, aging circadian rhythms have not previously been characterized in the context of immunosenescent cutaneous barrier defenses.

Interleukin 27 (IL-27), a member of the IL-12 family of heterodimeric cytokines, was recently implicated in cutaneous defense against Zika virus^9^. In response to skin injury, CD301b^+^ leukocytes are rapidly recruited to the wound site^10^, and produce IL-27 which subsequently potentiates wound closure and induces production of innate antiviral proteins (AVPs)^9, 11^. AVPs encompass several families, including oligoadenylate synthetase (OAS1, OAS2, OAS3), Myxovirus resistance proteins (MX1 and MX2), and interferon-induced transmembrane (*IFITM*) family proteins^1^. Circadian rhythms have been implicated in interferon-stimulated gene responses in several tissues, including skin and lung^12, 13^. However, it is unclear if skin barrier antiviral function is influenced by circadian rhythms.

In this study, we discovered that circadian factors Bmal1 and Clock decrease in aged skin. We also found that circadian dysregulation impairs cutaneous AVP expression via epidermal keratinocyte-autonomous and leukocyte-derived cytokine-mediated processes. Our studies show that murine cutaneous AVPs are regulated by Bmal1 and Clock in intact and wound states. We found that circadian-mutant and aged mice exhibited reduced IL-27 expression in skin wounds. Additionally, IL-27, along with type I interferon signaling, is required for time-of-day dependent circadian-regulation of wound-induced AVP production. We demonstrate that genetic loss-of-function of circadian factors sensitized skin and keratinocytes to HSV1 infection, whereas agents that increase circadian rhythm amplitudes in other tissues, such as SR8278 and nobiletin^14^, enhanced cutaneous circadian rhythms and reduced HSV1 viral burden in human keratinocytes and epidermal explants. Finally, we demonstrate that these circadian agents have antiviral effect in aging murine skin and human skin cells.

## RESULTS

### Aging skin displays a decline in circadian rhythm and antiviral immune function

The link between aging, cutaneous circadian rhythms, and barrier defense is unclear. To address this, we first investigated expression of circadian factors in murine skin of varying ages. We found that *Bmal1, Clock,* and *Per2* were downregulated in aged (greater than 12 months) skin compared to young (approximately 3-6 months) skin (**Figure 1a**). Serial passaging of primary human keratinocytes, which acts as a surrogate for human skin aging^15^, showed that circadian transcriptional activity decreased with increasing passage numbers (**Figure 1b**). This is also corroborated in an existing human skin dataset, where ARNTL(BMAL1) appears to peak in middle age before declining in expression^16^.

**Figure 1:**
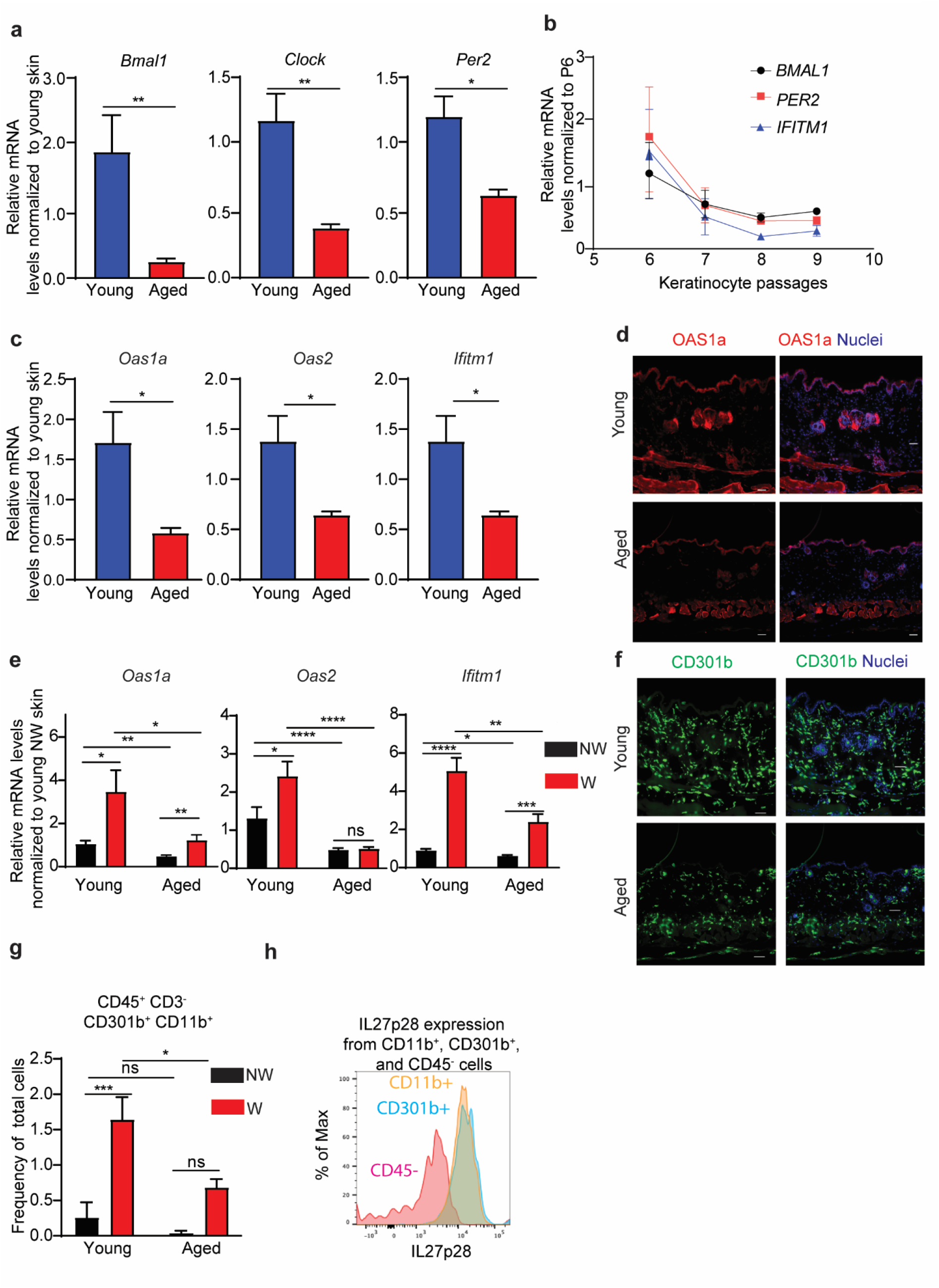
Aging skin exhibits diminished circadian, AVP, and IL-27 transcription. **a)** qPCR of *Bmal1, Clock,* and *Per2* in aged (n=14-16 1-year old) and young (n=32-34 1-month old) male murine skin. Graphs represent averages of relative mRNA **±** SEM with GAPDH used for internal control. P-values were obtained via Student’s T test. **b)** qPCR of *BMAL1, PER2* and *IFITM1* in human primary keratinocytes over serial passaging (n=2-3 donors per passage). Graphs represent averages of relative mRNA **±** SEM with GAPDH used for internal control. **c)** qRT-PCR of *Oas1, Oas2,* and *Ifitm1* in aged and young murine back skins as described in (a). **d)** Immunostaining for OAS1a [orange], nuclei [blue] in aged and young unwounded skin. Bar=25μm. **e)** qRT-PCR of AVP in young and old skin 24-hour post-wounding as described in (a). **f)** Immunostaining for CD301b [green], nuclei [blue] in aged and young unwounded skin. Bar=25μm. **g)** Flow cytometry showing reduced percentages of CD301b+ cells in aged skin compared to young skin (n= 4 mice per group). P-values obtained via Student’s T test. **h)** Histogram displays IL-27 production from CD11b+ (yellow) and CD301b+ (blue) cells compared to CD45-(red). Flow cytometry gating strategy included in Supplemental Figure S1.

We next investigated whether aging skin exhibits deficiency in antiviral protein (AVP) production. Using quantitative RT-PCR and immunofluorescence, we found that mRNA and protein levels of AVPs (*Oas1a, Oas2, Ifitm1*) were significantly reduced in aged murine skin compared to young skin (**Figure 1c-d**). Barrier disruption triggers an AVP response ^9, 11, 17^ in young skin. We observed that wounding significantly elevated AVP induction at 24 hours post wound in young and old skin; however, the magnitude of induction is significantly higher in wounds of young skin compared to that of aged skin (**Figure 1e**).

AVPs are induced in the skin wound microenvironment by IL-27, a cytokine produced by CD301b^+^ leukocytes of the skin ^9, 11^. To address if aging impacts CD301b^+^ signaling, we examined murine skin across ages for the presence of CD301b^+^ cells via immunostaining and found that CD301b^+^ cells were reduced in the skin of aged compared to younger mice (**Figure 1f**). Flow cytometry analysis revealed that wounded aged skin had a decreased influx of CD301b^+^ cells and expression of IL-27 (**Figure 1g-h**) (Gating strategy shown in **Supplemental Figure S1**). IL-27 works in concert with type I interferon in inhibiting Zika virus infection^9^. To test if type I interferon signaling is also affected by aging, we performed quantitative RT-PCR comparing the expression levels of type I interferons and their receptor, *Ifnar1,* in wounds of young and aged mice. We found no differences in *Ifna2*, *Ifnβ, Ifna4,* and *Ifna11* between old and young skin wounds collected 24 hours post-wounding (**Supplemental Figure S2**). This data supported a link between aging, cutaneous rhythms, and antiviral barrier defense. Next, we asked if circadian decline could mechanistically be responsible for this immunosenescent barrier defense.

### Cutaneous circadian rhythms regulate antiviral proteins

To test the circadian-innate immunity link in the context of skin, we queried published microarray gene expression data of murine skin harvested every 4 hours^18^. We scaled and clustered the expression of AVPs across Zeitgeber (standardized time-of-day) time-points within a 24-hour span using an additional dataset ^18^ and separated the genes into 5 clusters of distinct expression profiles. We found a variety of antiviral genes, including *Oas1a/g,* whose pattern of expression coincided with *Bmal1(Arntl*) in murine skin (**Figure 2a**). We then used a CIRCOS plot and determined a broader time-of-day regulation of antiviral immune genes in the skin (**Supplemental Figure S3**), which was supported by findings in baboon skin (**Figure 2b**) ^19^ where antiviral genes had rhythmic expression.

**Figure 2:**
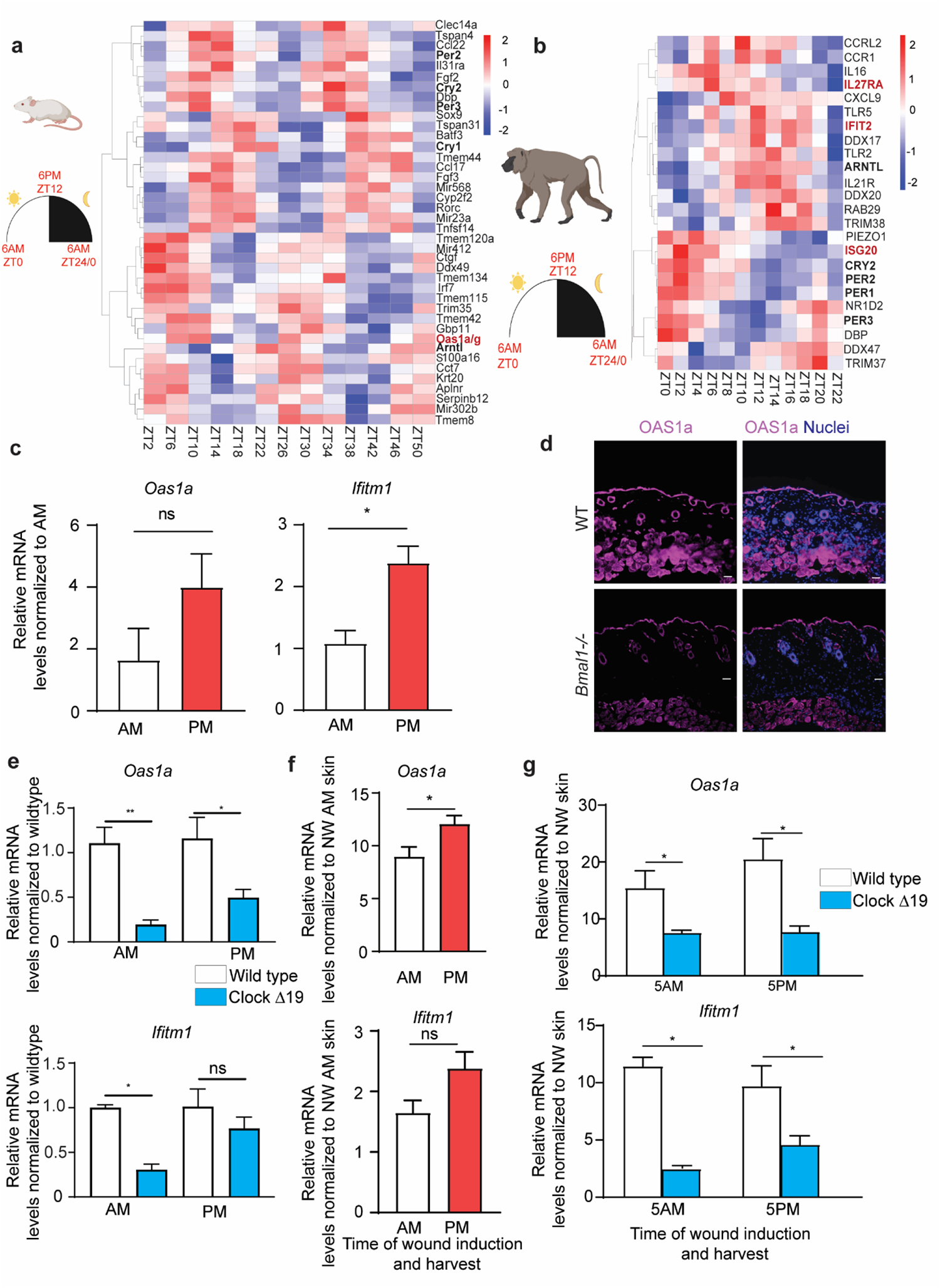
Circadian rhythm transcriptional networks include antiviral genes in mammalian skin. **a-b)** Heatmaps showing rhythmic expression of circadian factors and AVP genes in (a) murine skins (GSE38625) and (b) non-human primate baboon skins (GSE98965). **c)** qPCR of *Oas1* and *Ifitm1* in C57BL/6 murine belly skin harvested at 8AM or 8PM (n=4 mice per time-point). Graphs represent averages of relative mRNA **±** SEM with GAPDH used for internal control. P-values were obtained via Student’s T Test. **d)** Immunostaining for OAS1a [purple] in WT and *Bmal1^-/-^* skin. Nuclei [blue]. Bar=25μm. **e)** qRT-PCR of *Oas1a* and *Ifitm1* in intact skin of ClockΔ19 mice and BALB/C WT littermates at the indicated time (n=2-3 mice in each group with technical triplicates) as described in c). **f)** qPCR of *Oas1* and *Ifitm1* in skin wounds of C57BL/6 inflicted at times indicated and harvested 24 hours later (n=4 mice in each group) as described in c). *Ifitm1* p-value =0.0725. **g)** qRT-PCR of *Oas1a* and *Ifitm1* in skin wounds of ClockΔ19 mice and BALB/C WT littermates inflicted at the indicated time and harvested 24 hours later (n=2-3 mice in each group with technical triplicates) as described in c).

To validate this computational data and test if these AVP fluctuations are linked to circadian factors, we harvested belly skin from C57BL/6J mice at 8AM and 8PM time-points. We found that the basal expression of AVPs varied in the skin, as shown by qRT-PCR (**Figure 2c**). In agreement with the circadian AVP transcriptional data, Oas1a immunofluorescence staining was reduced in *Bmal1^-/-^* skin compared to wild type skin (**Figure 2d**). This was also supported by qPCR in the skin of circadian-deficient ClockΔ19 mutant mice ^20^ where circadian mutants expressed less AVP than their wild type littermates (**Figure 2e).**

To further examine the link between clock genes and AVP in the context of barrier disruption, we first compared AVP induction between wild type skin wounds collected at 8AM and 8PM, 24 hours post-wounding. *Oas1a* was significantly higher in 8PM wounds than that of 8AM wounds (**Figure 2f**); *Ifitm1* exhibited a similar trend, though did not reach significance. Compared to wild type counterparts, ClockΔ19 mutant skin exhibited a significant decrease of wound-induced AVP production across multiple time-points (**Figure 2g**). This supported a link between circadian rhythms and antiviral proteins of the skin.

### Circadian regulation of AVP-induction requires IL-27 and type I interferon signaling

Cytokine production and leukocyte trafficking are well-characterized immune phenotypes with a circadian level control^21^. We examined IL-27 expression in *Bmal1^-/-^*and *Bmal1*^+/-^ mouse skin, aged around 1 month, in an existing dataset ^18^. We noted decreased expression of IL-27 along with other interferon-responsive antiviral genes such as *Oas1a*, *Ifitm1 Ifitm7* and *Ifit3b* for intact *Bmal1^-/-^* skin (**Figure 3a**). Immunostaining revealed that numbers of CD301b^+^ cells were reduced for intact *Bmal1^-/-^*skin compared to wild type counterparts (**Figure 3b**). Flow cytometry analysis confirmed the decrease of CD301b^+^ cells in *Bmal1^-/-^*skin wounds and a reduced median fluorescence intensity (MFI) of IL-27p28 in these cells (**Figure 3c-d, Supplemental Figure S4**). To determine whether IL-27 is required for circadian-regulation of AVP induction, we used a Cre-loxP mouse model to ablate *Il27p28* in Lysozyme M-expressing (LysM-Cre) myeloid cells including CD301b^+^ cells^11^. We observed that deletion of IL-27 in myeloid cells markedly diminished the time-of-day response of wound-induction of AVPs (**Figure 3e**).

**Figure 3:**
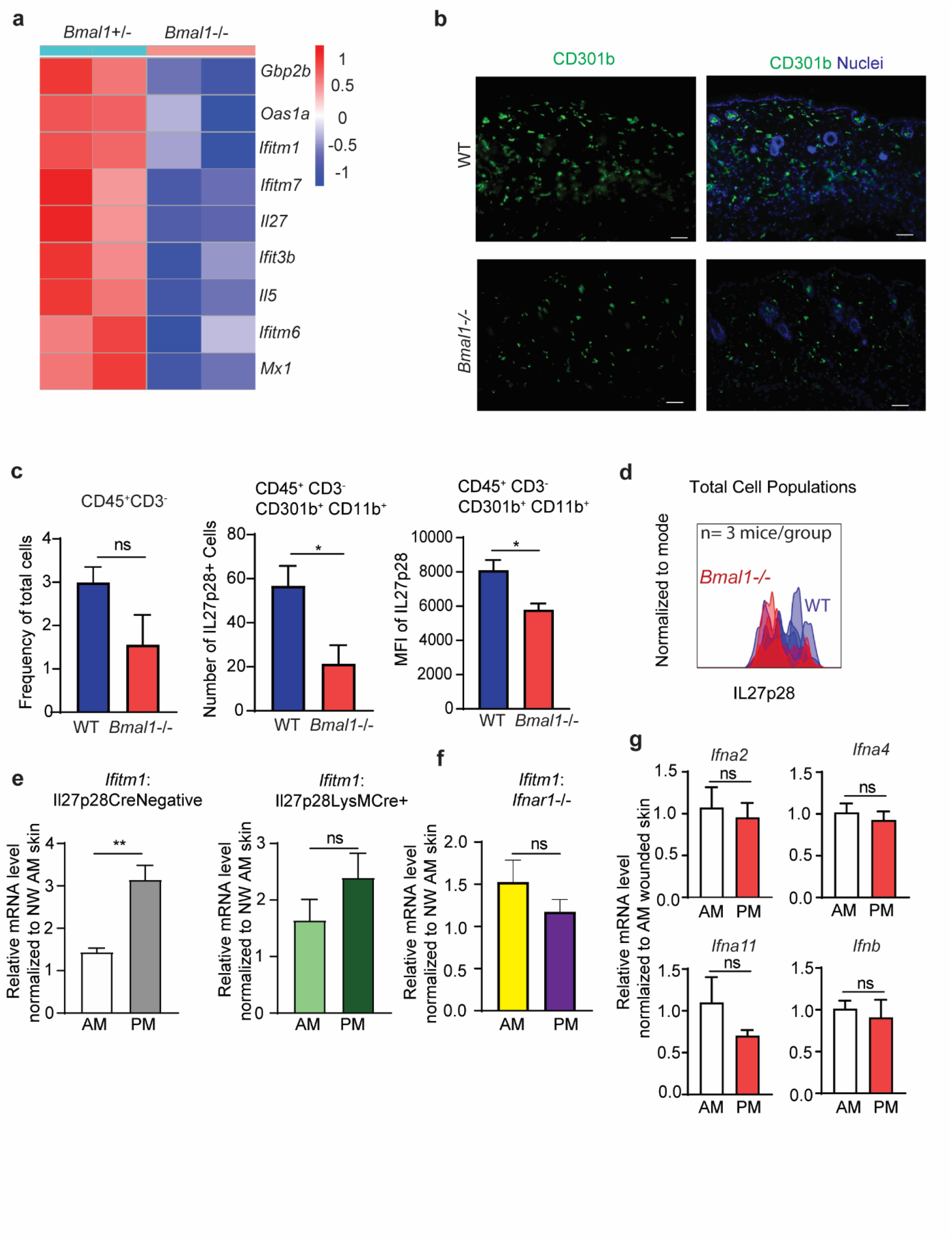
Circadian rhythms of wound-induced antiviral proteins require cytokine signaling. **a)** Heatmap of microarray expression of *Bmal1^-/-^* compared to heterozygotes intact murine skin as measured at Zeitgeber time 22 (GSE38625). **b)** Immunostaining for CD301b [green] and nuclei [blue] in WT and *Bmal1^-/-^* intact skin. Bar=25μm. **c)** Flow cytometry of CD301b+ cells and MFI of Il27p28 in *Bmal1^-/-^* skin and WT skin (n= 4 mice per group). **d)** Histogram displays IL27p28 expression in *Bmal1*^-/-^ (red) and WT mouse skin (Blue). Gating strategy is shown in **Supplementary Figure S4**. **e)** qRT-PCR of *Ifitm1* in skin wounds of WT or *LysM-*Cre.*IL27p28^fl/fl^*(n= 3 mice per group). **f)** qRT-PCR of *Ifitm1* in skin wounds of *Ifnar1^-/-^* mice (n=5-6 mice). **g)** qRT-PCR of type I interferons in WT C57BL6 mice wounded at 8AM or 8PM (n=4 mice per time point) as in e) above. Graphs represent averages of relative mRNA **±** SEM with GAPDH used for internal control. P-values were obtained via Student’s T Test.

To test if type I interferon has a circadian wound effect, we wounded *Ifnar1^-/-^* mice at 8AM and 8PM. We found that *Ifnar1* loss blunted significance of temporal variation in *Ifitm1* expression (**Figure 3f**). However, transcription of type I interferons, including *Ifna2*, *Ifna4*, *Ifna11*, and *Ifnb*, did not change with respect to time-of-day when measured 24 hours post-wounding (**Figure 3g**). This data supported a leukocyte mechanism of action for circadian-AVP function. Next, we asked if circadian-AVP regulation also operates at a keratinocyte-intrinsic manner.

### Keratinocyte-autonomous circadian rhythm regulates antiviral protein transcription and cutaneous defense against herpes simplex virus 1 infection

Cell-autonomous immune defects are present in circadian-deficient fibroblast cultures with respect to viral infection^6, 7^. We asked whether keratinocyte-autonomous circadian rhythms contribute to observed AVP regulation. To address this question, we synchronized circadian clocks of primary human epidermal keratinocyte cultures via an overnight incubation with omission of growth factor supplements. As expected, clock synchronization induced an oscillatory pattern of *BMAL1* expression (**Figure 4a**), coinciding with the oscillation of *OAS1, OAS2*, and *MX1* antiviral genes that approximate a cosinor sine model (**Supplemental Figure S5**). To establish a direct link between circadian factors and AVP expression, we performed siRNA-mediated knockdown of *CLOCK (siCLOCK)* in primary keratinocyte cultures. By qRT-PCR, we found that gene silencing of *CLOCK,* as confirmed by RT-PCR, significantly reduced expression of *IFITM1* (**Figure 4b**). Furthermore, this effect had direct effects on viral replication; when primary keratinocytes with siBMAL1 and siCLOCK were infected with herpes simplex virus type 1 (HSV1), they produced more virus than nonsilencing siControl as measured by viral PCR (**Figure 4c**). This was corroborated via immunofluorescence using an immortalized keratinocyte model, *NTert*s, where circadian disruption was associated with significantly increased HSV1 antigen levels (**Figure 4d-f).**

**Figure 4:**
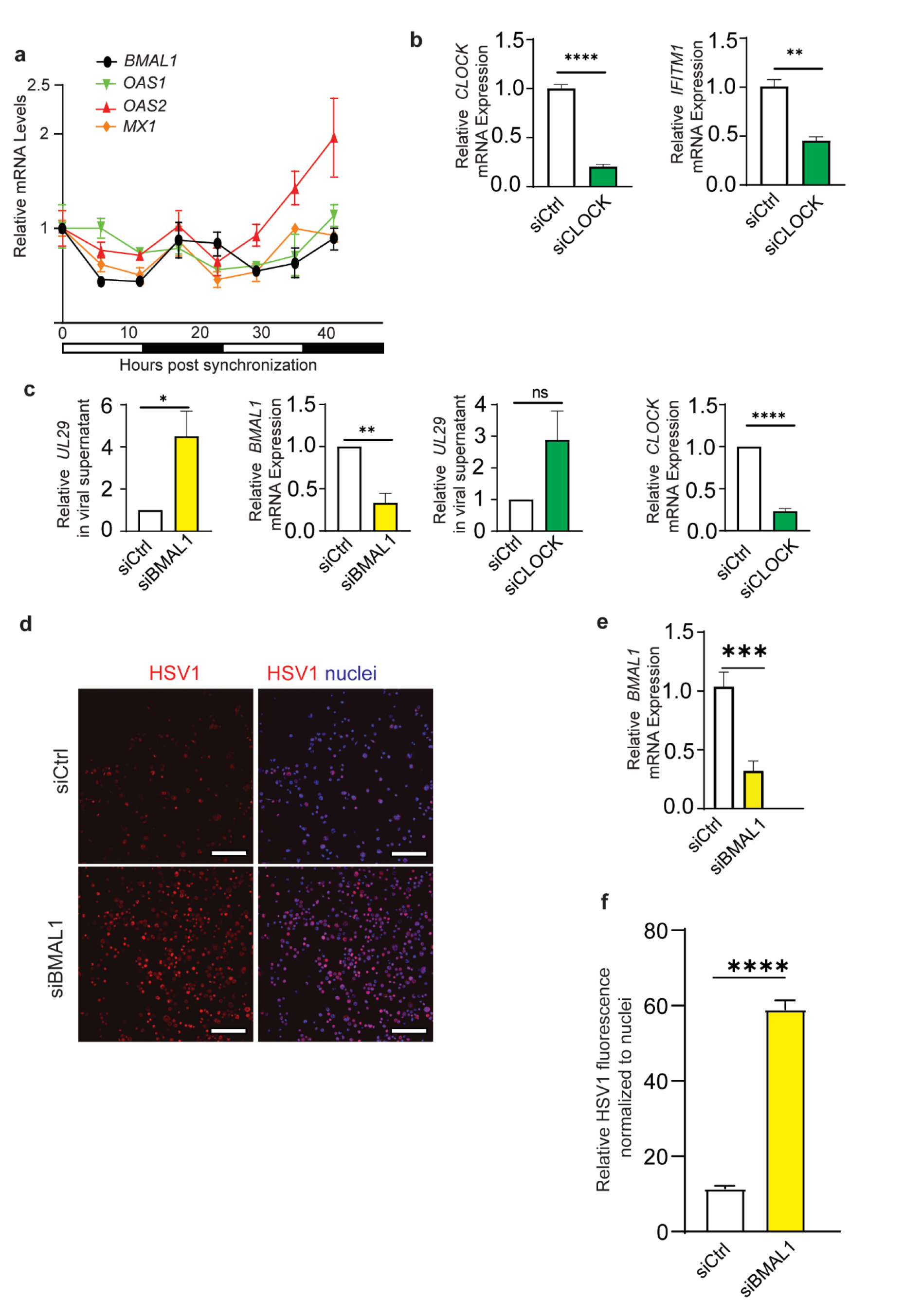
Keratinocyte autonomous circadian rhythm regulates antiviral activity. **a)** qRT-PCR of *BMAL1, OAS1, OAS2,* and *MX1* in human primary keratinocytes synchronized via growth factor starvation and harvested every 6 hours (representative of three independent experiments). Graphs represent relative mRNA **±** SEM with GAPDH used for internal control and relative to that of 0-hour time-point. **b)** qRT-PCR of *CLOCK* and *IFITM1* in human primary keratinocytes that received small interfering RNA, either non-silencing control (siCtrl) or specific for CLOCK (siCLOCK) (n=3 technical triplicates, representative of three independent experiments). Graphs represent averages of relative mRNA **±** SEM with GAPDH used for internal control. **c)** qPCR of HSV1 gene UL29 in supernatant of primary keratinocytes that received transfection of either siCtrl, siBMAL1, or siCLOCK 24 hours prior to infection with HSV1 (MOI 0.01). Knockdown efficacy was shown by qRT-PCR. Graphs represent averages of either relative DNA or mRNA **±** SEM with GAPDH used for internal control. P-values were obtained via Students T test. n=3 primary keratinocyte donors. **D)** Immunofluorescence of HSV1 (MOI 0.01) in human *NTERT* Keratinocytes transduced with non-silencing siCtrl or siBMAL1. Bar= 160um. **E)** Knockdown efficacy of BMAL1 in human *NTERT* shown by qRT-PCR. **F)** ImageJ (Fiji) quantification of relative viral immunofluorescence normalized to nuclear staining. Graphs represent averages of relative immunofluorescence **±** SEM (n=3 samples). P-values obtained using (a-b) via Student’s T test and (c-d) ANOVA with multiple comparisons.

### Circadian enhancement leads to decreased HSV1 infection in the skin

We hypothesized that enhancement of circadian function increases antiviral immunity of the skin. We expressed a *Bmal1* promoter-driven luciferase reporter^22^ in *NTert* keratinocytes and validated that *Bmal1*-reporter expression is regulated in a circadian-dependent manner (**Figure 5a**). Using the *Bmal1*-reporter system, we found that treatment of 10 µM SR8278, a small molecule REV-ERB antagonist previously shown to increase circadian rhythms in non-skin tissues^23^, enhanced the amplitude of rhythmic BMAL1 activity (**Figure 5a-b**).

**Figure 5:**
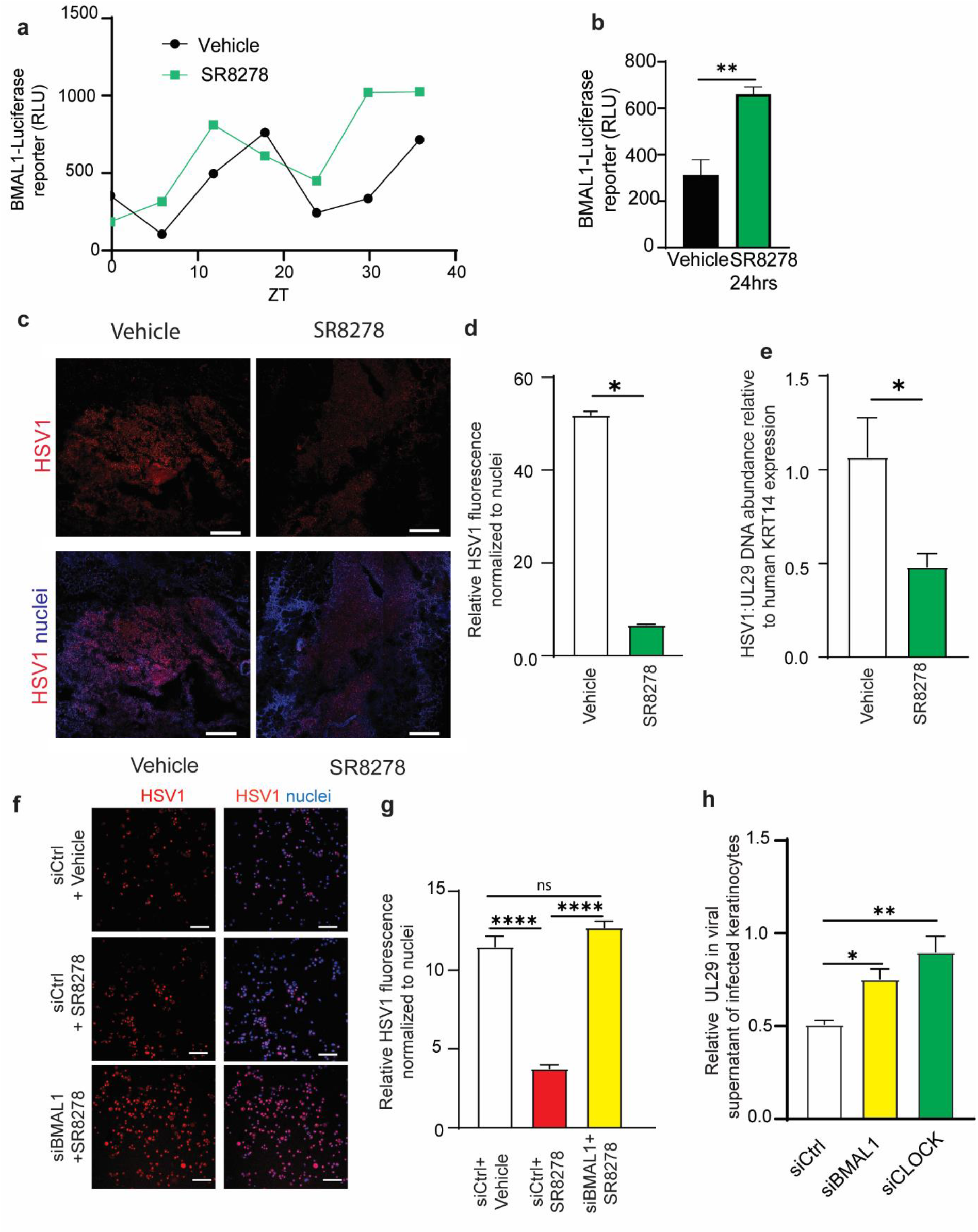
Pharmacological augmentation of circadian rhythm reduces herpes simplex viral infection in a BMAL1/CLOCK-dependent manner. **a-b)** *Bmal1*-luciferase reporter assay. Human *NTERT* keratinocytes were transduced with *Bmal1*-luciferase reporter construct and growth supplement starved overnight before treatment with vehicle or 10 µM SR8278. Cells were harvested (a) every 6 hours and (b) at 24 hours for luminescence measurements (n=4). **c)** Immunostaining for HSV1 antigen in human epidermal explants infected with HSV1 and treated with vehicle or 10 µM SR8278. Bar=500um. **d)** ImageJ (Fiji) quantification of viral immunofluorescence normalized to nuclear staining. **e)** qPCR of HSV1 viral gene UL29 relative to human KRT14 in epidermal skin infection. **f)** Immunostaining for HSV1 antigen in human *NTERT* keratinocytes (MOI 0.01) transduced with siCtrl or siBMAL1 and supplemented with vehicle or 10 µM SR8278. Bar = 160um. **g)** ImageJ (Fiji) quantification of relative viral immunofluorescence normalized to nuclear staining. Quantification of non-silencing control (SiControl, vehicle) is same data as used in Figure 4e. **h)** qPCR of HSV1 viral gene UL29 in supernatant of infected primary keratinocytes (MOI 0.01) transfected with siRNA against BMAL1 or CLOCK. Graphs represent averages of relative DNA or viral immunofluorescence normalized to nuclear staining **±** SEM. P-values obtained using Student’s T test or ANOVA with multiple comparisons.

To test if circadian augmentation has an antiviral effect, we utilized surgically discarded human skin samples. We separated human epidermis from dermis, infected the epidermis *ex vivo* with HSV1 in an air-liquid interface culture system and treated the epidermal explant culture with either vehicle or 10 uM SR8278 for 24 hours. By immunofluorescent staining and quantification, we observed that SR8278 treatment significantly reduced HSV1 antigen expression in the epidermis (**Figure 5c-d**). We confirmed this reduction via qPCR for HSV1 UL29 gene with human K14 gene used for internal control (**Figure 5e**). We then asked whether the antiviral effect of SR8278 was dependent on circadian and antiviral factors. We performed siRNA-mediated gene silencing of BMAL1 and CLOCK and found that the antiviral effect of SR8278 was significantly lessened in cells transfected with siBMAL1 and siCLOCK compared to non-silencing control cells (**Figure 5f-h).** Interestingly, when we suppressed antiviral proteins OAS1 and IFITM1 via siRNA in *NTERT* keratinocytes, SR8278’s effect was also lessened (**Supplemental Figure S6**) **s**uggesting SR8278’s antiviral effect is both circadian and antiviral protein dependent.

We subsequently examined if other circadian augmenting compounds would have antiviral effects. Nobiletin is an antioxidant flavonoid with multiple pharmacological effects, including antioxidant properties^24^, as well as an ROR agonist that potentiates circadian rhythms ^14^. Using the *NTert Bmal1*-luciferase cell line, we confirmed that nobiletin increased *Bmal1*-reporter expression in keratinocytes (**Supplemental Figure S7a**). Treatment of nobiletin decreased HSV1 infection of human epidermal explants (**Supplemental Figure S7b-d**), suggesting that our findings are not drug-specific, but rather related to the circadian rhythm effect.

### Viral infections in aging skin are reduced by circadian augmentation

Aging skin is subject to immunosenescence and is known to be susceptible to viral infections. Thus, we examined if circadian modulation has antiviral activity in aging skin. We found that 1 year-aged *Bmal1^+/-^*mutant aging skin, which display premature aging, also show deficiency of antiviral protein transcription in the wound environment (**Figure 6a**). Using a mouse epidermal explant infection model, we found via qPCR that HSV1 levels were higher in aged skin than that of younger skin **(Figure 6b).** Treatment of circadian drug SR8278 reduced HSV1 viral DNA load in aged skin by roughly 50% as measured by qPCR **(Figure 6c)**. We then returned to our passaging model of human keratinocytes as a pseudoaging infection of human skin. We found that human primary keratinocytes at P8 produce more virus than P2 keratinocytes (**Figure 6d),** and that this increased viral replication could be suppressed by treatment of SR8278 **(Figure 6e)**. These data indicate that circadian augmenting agents have antiviral effects on aging skin.

**Figure 6:**
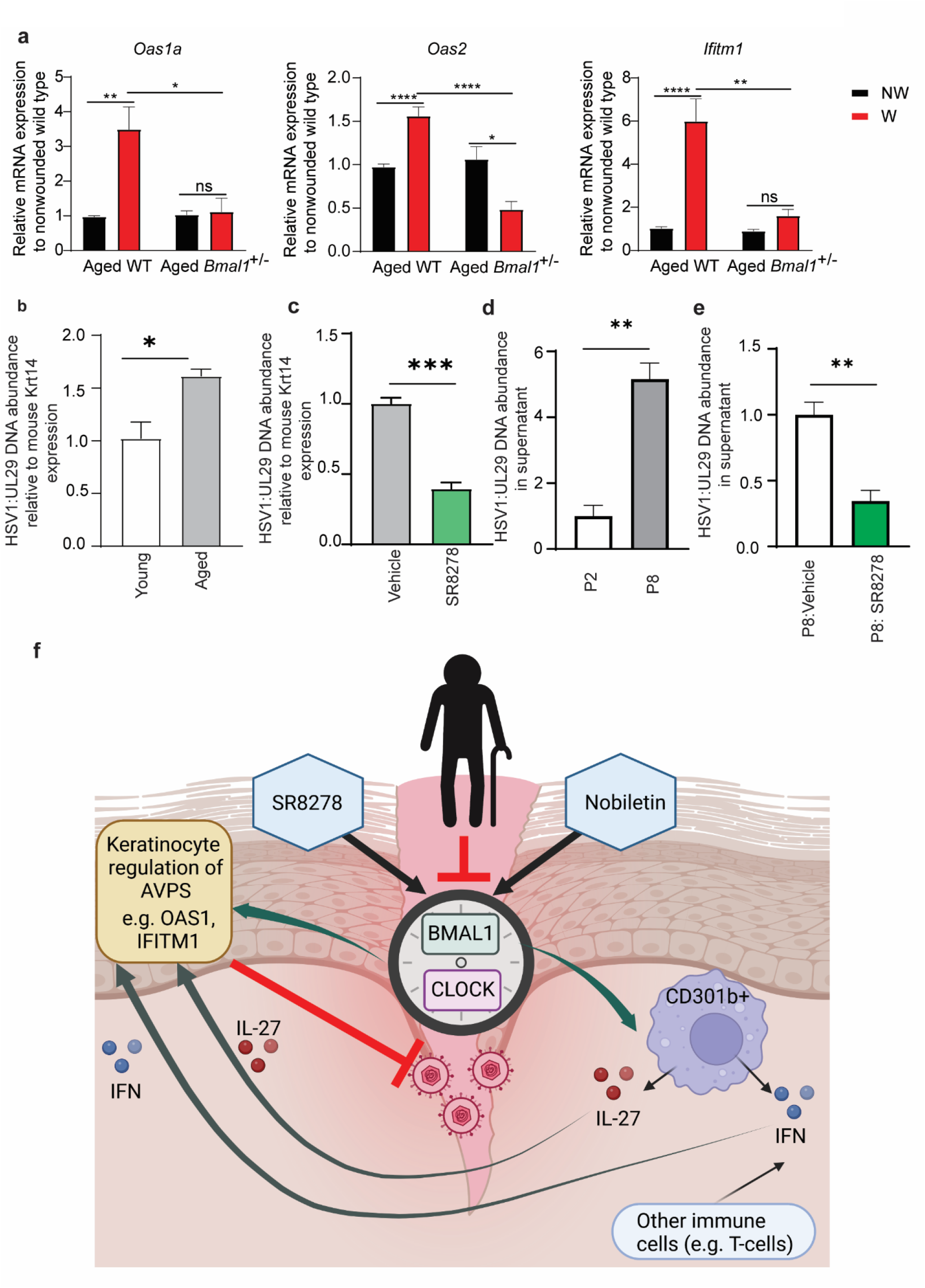
Antiviral immune decline of aging skin can be rescued by a circadian enhancer treatment. **a)**. qRT-PCR of Oas1a, Oas2, and Ifitm1 in Bmal1^+/-^ and wild type skin that was wounded and collected 24 hours later (n=3-6 mice per genotype). **b)** qPCR of viral UL29 relative to murine Krt14 in HSV1 infection of aging mouse (>365 days) and young mouse (2-6 months) epidermis (n=3 skin explants of each age). **c)** qPCR of viral UL29 relative to murine Krt14 in HSV1 infection of aging mouse (>365 days) epidermis treated with vehicle or 10uM SR8278 (n=3 skin explants per condition). **d-e)** qPCR of HSV1 UL29 in supernatants of infected P2 and P8 keratinocytes treated with vehicle or 10uM SR8278 (MOI 0.01). Graphs represent averages of relative HSV1 DNA to mouse Krt14 ± SEM. P values obtained using Student’s T test n=3 primary donors of each age. **f)** Working model of aging-associated decline of cutaneous innate antiviral immunity. Circadian rhythm factors BMAL1 and CLOCK regulate expression of cutaneous antiviral proteins in a keratinocyte-autonomous manner and also via a leukocyte-mediated process, where IL-27 conveys a time-of-day response and type I interferon signaling ensures a robust antiviral immunity. Aging-associated circadian decline decreases cutaneous antiviral immunity, while pharmacological means of circadian enhancement increases it.

## DISCUSSION

Our data reveal a pharmacologically tractable model of age-mediated circadian regulation of antiviral immunity of the skin. Skin aging leads to decline of circadian function in the skin, compromising epidermal keratinocyte and dermal leukocyte antiviral responses. Pharmacological agents that potentiate skin cell circadian amplitude improve immunity via antiviral protein effect. Our data delineate new mechanisms responsible for the immunosenescent decline of antiviral immunity in aging skin, underscoring the circadian pathway as a new therapeutic target for enhancing aging skin barrier function.

Previous studies have shown that interferon stimulated genes are affected by skin circadian rhythms through a TLR7-dependent stimulation^12^. We have demonstrated that type I interferon signaling is required for activation of antiviral immune responses but may not convey time-of-day responses. However, type I interferon signaling is required for maximum AVP expression irrespective of time-of-day, suggesting that the interferon pathway as the primary regulator of AVPs is subject to regulation by circadian factors. In this regard, STAT1 and STAT3, transcription factors involved in interferon response, display time-of day-response^25^, and could be an indirect regulatory step between circadian transcription factors and antiviral proteins. Circadian factors may directly regulate AVPs by binding to the E-box consensus elements which are present in gene promoters of antiviral genes, such as *Oas1*, *Oas2*, and *Ifitm1*. Such possibilities may be explored via skin cell-specific ChIP-Seq for BMAL1 and CLOCK^26^. These experiments may also help explain why certain AVPs in primate skin follow distinct temporal expression (**Figure 2B**) as AVPs may experience differential BMAL1/CLOCK binding efficacies in the skin.

Our data show that CD301b^+^ leukocyte-derived IL-27 is important for the time-of-day-dependent response of AVP expression. It will be important to determine whether the decreased dermal infiltration of CD301b^+^ leukocytes in aged skin and *Bmal1*^-/-^ skin is a result of central or local circadian decline, and if rescuing this defect can restore circadian AVP functionality. Knowing specifically how the circadian rhythm acts on circulating immune cells and skin resident cells will provide better insights into how to leverage circadian rhythms for improving cutaneous tissue regeneration and defense against infection in aging skin. This information may also prove useful in treatment of skin infections that resist traditional interferon induced immunity, such as monkey pox ^27^.

We focused on a common skin pathogen HSV1, but intriguingly, different viruses may interplay with circadian rhythms in distinct fashions. For example, respiratory syncytial and vesicular stomatitis virus replication rates were found to have opposite responses to circadian deletion^7^, possibly due to how these viruses differ in viral entry and replication machinery. Other work has shown that while the circadian rhythm impacts response to HSV-2 in murine skin, it has minimal impact on survival^28^. Our data show that siRNA-mediated gene silencing of *BMAL1*/*CLOCK* sensitized human keratinocytes to HSV1 infection and diminished protective effects conferred by pharmacological circadian enhancers (SR8728 and nobiletin). In agreement with our data, *Bmal1^-/-^* mice exhibit greater HSV1 viral replication than wild type counterparts^6^, which is in part attributed to aspects of host cell-virus interaction, such as intracellular trafficking and chromatin assembly. Paradoxically, BMAL1 and CLOCK are found to be hijacked by viral proteins to support viral replication^29^. How to best balance the antiviral and pro-viral activity of circadian factors requires further study and should incorporate more skin-trophic viruses that have pandemic level infectivity risks^30^.

In summary, we demonstrate a novel mechanism of aging-associated skin infection risk. Age leads to circadian suppression in the skin, ultimately triggering a reduced antiviral barrier function in a BMAL1/CLOCK-dependent manner. Circadian pharmacological agents are able to rescue age related viral susceptibility in the skin, suggesting a novel therapeutic pathway for combating immunosenescence. Our findings have potentially wide implications for aging skin and may lead to new treatment strategies for prevention of cutaneous infection, wound care, and overall skin health in aging populations.

## MATERIALS AND METHODS

### *In vivo* wounding experiments

C57BL/6 wild type, *Ifnar1^-/-^*, B6.129-Arntltm1Bra/J (*Bmal1^-/-^*), ClockΔ19 mice ^20^ and LysM*-*Cre mice^20, 31^ were obtained from Jackson laboratories (Bar Harbor, ME). *IL27p28^fl/fl^* mice were gifts from Zhinan Yin (Biomedical Translational Institute, Jinan University) and Li Fan Lu (University of California San Diego). All mice were maintained under a specific pathogen-free environment. After anesthesia of the mice, 3-mm punch wounds were made on the back of each mouse at distinct times of day and collected 24 hours post-wounding. Tissue from non-wounded and wounded skin was dissected from each mouse, lysed in Trizol reagent (ThermoFisher, Waltham, MA), and kept at -80°C for RNA extraction, placed in OCT for immunofluorescence, or immunostained for flow cytometry. Information regarding RNA extraction, quantitative real-time PCR, immunofluorescence, or flow cytometry approaches can be found in Supplementary Methods. For aged/young skin studies, wounds were inflicted between 8:00 AM and 9:00 AM. Young mice were between 3-6 months, and elderly mice were greater than a year. Male and female mice were used in this study.

### Keratinocyte cell culture

Human primary keratinocytes were purchased from Thermo-Fisher Scientific. Cells were grown in a 37°C incubator in serum free Epi-Life cell culture medium (Gibco, Waltham, MA) supplemented with Epi-Life Defined Growth Supplement (EDGS) containing 0.05mM Ca^2+^. *N/Tert 2G* keratinocytes were a gift from the lab of Johann Gudjonsson of University of Michigan and cultured in Keratinocyte SFM media supplemented with EGF and BPE (Gibco) prior to use for luciferase and infection studies. Complete keratinocyte methods, including circadian synchronization, siRNA experiments, and infections can be found in supplementary methods.

### RNA-Seq gene expression and microarray gene expression data

Publicly available non-human primate RNA-Seq gene expression data was downloaded from the Gene Expression Omnibus (GSE98965). The data was provided as a normalized expression matrix as calculated by Mure et al. ^19^. Publicly available microarray gene expression data from Geyfman et al. ^18^ was downloaded from the Gene Expression Omnibus ^32^ (GSE38625). Further computational analysis methods can be found in Supplementary Methods.

### Herpes simplex virus staining and quantitative PCR

For full description of viral infection, please see Supplemental Methods. Briefly, human or murine epidermis maintained in keratinocyte growth media and infected with 10,000 focus-forming units per sample of herpes simplex virus type 1 (HSV1) strain NS in the presence of vehicle, nobiletin or SR8728 (Sigma) at 5-10 µM. The epidermis was then either placed in 4% formaldehyde to fix overnight and subsequently stained for HSV1 antigen or lysed for DNA extraction and viral quantification.

### Statistical analysis

All statistical tests were performed in GraphPad Prism. Throughout figures, data are shown as mean ± SEM, * =p ≤ 0.05, **= p ≤0.01, ***=p≤0.001, *\*\*\*\*=*p ≤ 0.0001. Diagrams and visual abstract were created using BioRender software.

### Study Approval

Animal procedures were performed in agreement with the recommendations in the Guide for the Care and Use of Laboratory Animals of the National Institutes of Health. Animal protocols were approved by Duke University’s Institutional Animal Care and Use Committee (Animal Welfare Assurance). Human tissue was used in accordance with Duke Institutional Review Board approval.

## ACKNOWLEDGEMENTS

This study was supported by funding from:

National Institutes of Health 5R01AI139207 to ASM and JYZ

National Institutes of Health 5R01AR073858 to JYZ

National Institutes of Health R01AI139425 to JC

We thank Helen Lazear and Drake Philip (University of North Carolina, at Chapel Hill) for providing Herpes Simplex Virus and insight and guidance on viral infections. We thank Drs. Russell Hall and Suephy Chen (Duke Dermatology), Stacy Horner and Andrew Alspaugh (Duke MGM) for comments, the lab of Johan Gudjonsson (University of Michigan) for N/Tert 2G keratinocytes, and Zhinan Yin and Li Fan Lu (UC San Diego) for providing *Il27p28^fl/fl^*mice. We also thank Jacob Benton, Paula Marriottoni, Xin Ling, Margaret Coates and Yingai Jin (Duke University) for their technical assistance and valuable comments.

## Author Contributions

Conceptualization: SK, VL, AM, DH, JZ

Methodology: SK, VL, JS, AM, JZ, JC, DC

Investigation: SK, VL, PK, MP, DC, AM, JZ, DW

Visualization: SK, VL

Funding acquisition: JZ, AM, JC

Project administration: JZ, AM, JC, KD

Supervision: JZ, AM, JC, KD

Writing – review & editing: everyone

## Data Availability and Material Sharing

All data needed to evaluate our conclusions are present in the paper and/or the Supplementary Materials. Previously published datasets including^18, 19^ are cited in this paper, and data is publicly available. The data and materials generated in this study can be provided by Jennifer Zhang pending scientific review and a completed material transfer agreement. Requests for the data and materials should be submitted to: Jennifer Zhang.

