## Supplemental Materials for "An Aging-Susceptible Circadian Rhythm Controls Cutaneous Antiviral Immunity"

**Supplementary Data includes:**

Supplementary Methods
Supplementary Figure Legends
Figs. S1 to S7
Tables S1
Supplementary References

### SUPPLEMENTARY MATERIALS
### Supplementary Methods:
#### Keratinocyte Transfections and Circadian Synchronization

Circadian synchronization was performed with growth supplement starvation. Starvation was performed after plating by replacing full media with supplement free media overnight, and then fed with supplement replete media. Cells were harvested at various time-points post-synchronization. Synchronization experiments were performed 2 independent times with at least 2 different keratinocyte donors. For passaging experiments, cells were trypsinized and an aliquot harvested for RNA extraction at time of passage. Multiple keratinocyte donors were pooled at each passage and run as technical duplicates.

*NTert* keratinocytes were transfected with the pAB-Puro-BluF (Brown et al., 2005) plasmid (Addgene, Watertown, MA) and validated for stable expression prior to growth supplement starvation overnight. Cells were treated with 5 uM of nobiletin or 10 uM SR8278 and collected at ZT0-48. Luminescence was measured using the Promega Dual-Glo Luciferase Kit and Glomax-Multi Jr Single Tube Luminometer (Promega, Madison, WI).

For gene silencing of primary keratinocytes, three siRNA sequences designed to target CLOCK, BMAL1, OAS1, or IFITM1 were combined for maximal knockdown (Dharmacon). siRNA were transfected into keratinocytes using Genmute Reagent (SignaGen, Rockville, MD) at 10 nM concentration. RNA was collected 48 hours after transfection to evaluate knockdown efficacy. Expression changes were evaluated relative to non-silencing siControl (Origene).

**Keratinocyte Circadian Oscillation**

For our circadian oscillation transcription data, we have modelled our data using Graphpad Prism software and the R studio package 'Psych' which includes a cosinor sine model that fits a circadian sine wave to data. Fit to a true circadian oscillation was evaluated using this data. Periodicity, Mesor, amplitude, and acrophase of genes were determined using Cosinor Software (Refinetti)

**RNA extraction and qRT-PCR**

RNA was extracted using the Direct-zol RNA Purification Kit (Zymo Research, Tustin, CA). cDNA was transcribed using the iScript cDNA synthesis kit (BioRad, Hercules, CA) and resulting cDNA was used for quantitative RT-PCR with SYBR Green Master Mix (ThermoFisher) on the StepOne Plus Real-Time PCR machine (Applied Biosystems, Foster City, CA). Primers used for amplification of targeted genes are shown in Supplementary Table 1. Fold change of gene expression was calculated and normalized to the housekeeping gene GAPDH and calculated using the $2^{(-\Delta\Delta Ct)}$ method (Livak and Schmittgen, 2001).

**Immunofluorescence**

Mouse skin was frozen in O.C.T. for sectioning. 10 μm sections were fixed in 4% paraformaldehyde, washed in PBS, and permeabilized in 0.1% Triton X-100 (10 minutes) and blocked in a blocking buffer containing 10% normal goat serum, 5% normal donkey serum 1% BSA, and 0.01% Triton X-100 (1 hour). The sections were then incubated overnight with primary antibodies (4 °C): rabbit polyclonal OAS1 (Millipore, Burlington, MA), CD301b (BioRad, Hercules CA), or rabbit IgG as a control (SouthernBio, Birmingham, AL), followed by washing and incubation with secondary antibodies (Cy3-conjugated secondary antibodies; Thermo

Scientific), washed in PBS containing 0.01% Triton X-100 and counterstained with Hoechst. Cells were imaged using the Olympus Cell Sens software or using a Zeiss 780 Confocal microscope. Fluorescence quantifications were performed using Fiji Software.

**Mouse skin cell isolation and flow cytometry**

24 hours post-wounding, skin wounds were harvested along with a 1-mm rim around the edge of the wound, minced, and pooled in 0.3% trypsin/0.1% glucose, 14.8 mM NaCl, 5.3 mM KCl (GNK) with 0.1% DNase at 4 °C overnight. Cell solutions were then incubated with monensin and brefeldin A prior to extracellular staining. Intracellular staining was achieved via fixation and permeabilization. Single-cell suspensions were washed and stained with the following antibodies: BV510-Live Dead, BV421-CD45, AF488-CD3, PE-CD301b, PE-Cy7-CD11b, APC-IL27p28 (BioLegend, San Diego CA). Flow cytometry was performed on a FACS Canto and was analyzed using FlowJo (Ashland, OR) Software.

**RNA-Seq gene expression**

Publicly available non-human primate RNA-Seq gene expression data was downloaded from the Gene Expression Omnibus (GSE98965). The data was provided as a normalized expression matrix as calculated by Mure et al. (Mure et al., 2018). Select genes that were identified as being significantly rhythmically expressed in the skin by their study had their expression log2 transformed, converted to z-scores, clustered using a correlation distance with complete linkage, and then plotted in a heatmap.

**Microarray gene expression data**

Publicly available microarray gene expression data from Geyfman et al. (Geyfman et al., 2012) was downloaded from the Gene Expression Omnibus (Edgar et al., 2002) (GSE38625). The raw data was normalized using the robust multi-array average (RMA) approach using the *affy* (Gautier et al., 2004) Bioconductor (Huber et al., 2015) package from the R statistical programming environment (Team, 2020). Select genes from the *Bmal1*$^{-/-}$ and *Bmal1*$^{+/-}$ samples were z-score transformed and displayed on a heatmap with the genes and samples being clustered by a correlation distance with complete linkage. Similarly, for the time-course samples, select AVPs were z-score transformed and plotted in a heatmap where the genes were clustered using a correlation distance with complete linkage. The genes were then split into 5 clusters based on distinct expression profiles and correlation clustering. A CIRCOS plot was used to show the expression profile of those gene clusters across the 13 time points. The ribbons are color coordinated to match the cluster definition from the heatmap. Ribbons are connected to time points where a majority of the genes within the cluster show expression above the mean. Ribbon width is proportional to the number of genes in each of the clusters. Similar to the Mure et al. study, we employed the meta2d integrative method from the *MetaCycle* R package to identify rhythmically expressed genes. Probe sets were considered rhythmically expressed if they had an FDR adjusted p-value <0.05.

**Herpes simplex virus staining and quantitative PCR**

For skin viral infection, human or murine skin explants were trimmed of fat, and epidermis and dermis separated in a 5mg/ml dispase solution overnight. Epidermis was subsequently maintained in keratinocyte growth media and infected with 10,000 focus-forming units per sample of herpes simplex virus type 1 (HSV1) strain NS in the presence of vehicle, nobiletin or SR8728 (Sigma) at 5-10 µM. The epidermis was then either placed in 4% formaldehyde to fix overnight or lysed for DNA extraction. Fixed tissues were stained as whole mount for HSV1 using Abcam antibody 9533 at a 1:60 dilution overnight and a 1:400 secondary antibody stain. For keratinocyte studies, Ntert 2G keratinocytes or primary keratinocytes were transfected with siRNA oligonucleotides specific for CLOCK or BMAL1 or non-silencing control at 10 nM. Cells were infected with HSV and treated with vehicle, 10 µM nobiletin or SR8278 for 24 hours. HSV1 stocks were grown in Vero cells and viral genomes quantified via PCR. For DNA level quantification in skin, epidermis was placed in DNA/RNA shield (Zymo) and DNA extracted before use in quantitative PCR. Human KRT14 was used as an internal control for the amount of host DNA present in each sample. Viral supernatant quantifications were used for cell infections and standardized to a viral standard curve of known viral quantity.

**Supplementary Table 1:** Primer Sequences

| **Primer** | **Forward Sequence** | **Reverse Sequence** |
|---|---|---|
| Human *GAPDH* | CAAGAGCACAAGAGGAAGAGAG | CTACATGGCAACTGTGAGGAG |
| Human *BMAL1* | CGGAGTCGATGGTTCAGTTT | CTTCCAGGACGTTGGCTAAA |
| Human *OAS1* | GTCTTCCTCAGTCCTCTCACG | AAGGCAGGCAGCACATCG |
| Human *CLOCK* | AGTGGTTCTGTGCCTCTTATC | CCAAGTCTACTGGACTGTCTTC |
| Human *OAS2* | TGAAGATGAGACCGTGAGGAAG | CCAGAAGATGCCAACACCAAC |
| Human *MX1* | GGACATCGCCACCACAGAGG | TCCGCACCACATCCACAACC |
| Human *KRT14* | CGTGACCAGTATGAGAAGATGG | CTCCTCTGTCTTGGTGAAGAAC |
| Mouse *Gapdh* | GCACAGTCAAGGCCGAGAAT | GCCTTCTCCATGGTGGTGAA |
| Mouse *Oas2* | CCGGGCCAGTGCACAAGTTAG | CGATGGCACCGAGGACACC |
| Mouse *Mx1* | GGGGAGGAAATAGAGAAAATGAT | GTTTACAAAGGGCTTGCTTGCT |
| Mouse *Oas1a* | GGAGGCGGTTGGCTGAAGAGG | GAACCACCGTCGGCACATCC |
| Mouse *Ifitm1* | CAGGAAGATGGTGGGTGATAC | CGTGAGGATGGTGAAGAACA |
| Mouse *Ifnar1* | ATAGCTGGGTCCTCAGGTAATA | TCCATTGCTGGTGGGAATAC |
| Mouse *Ifna2* | CTGCTGGCTGTGAGGAAATA | AGCAAGTTGACTGAGGAAGAC |
| Mouse *Krt14* | ACTCACTCGCTCACTTGCTCA | ATCTTGCTCTTCAGGTCCTC |
| HSV UL29 | CCATCATCTCCTCGCTTAGG | AGCTGCAGATCGAGGACTG |

**Supplemental Figure S1: Stepwise gating strategy for Figure 1g**

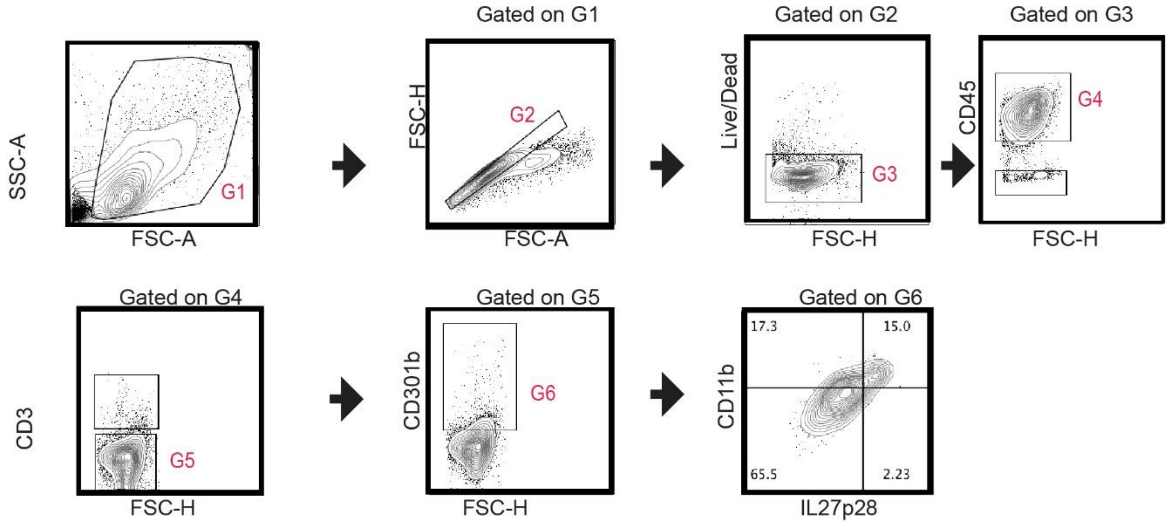


**Supplementary Figure S1:** Gating strategy for aging skin wounds**.** Aging skin was wounded, and flow cytometry used to gate to IL-27 producing cells. Cells were pre-gated on live single cells that were CD45+, CD3+, CD301b+, and CD11b+.

**Supplemental Figure S2: Expression of type I interferons in young and old wounds**

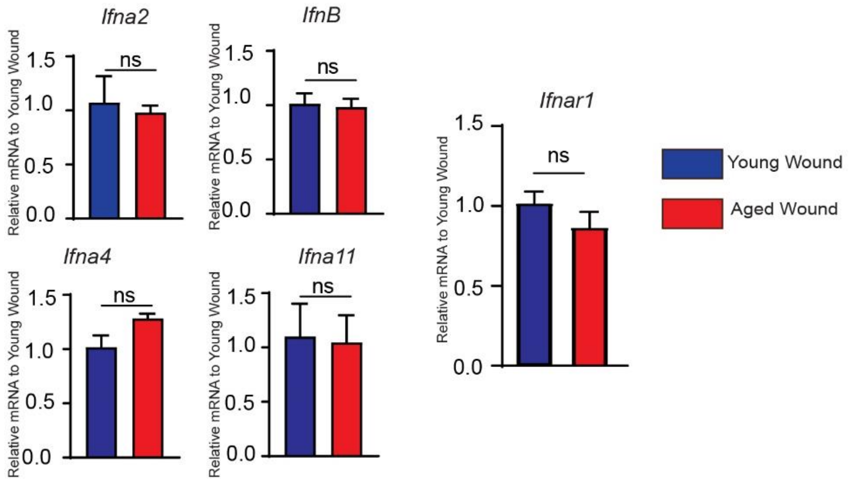


**Supplementary Figure S2:** Type 1 interferon expression in young and aged skins. qRT-PCR of *Ifna2* and *Ifnar1* in aging (over a year) and young (2-3 month) mice in wounded tissue collected 24 hours post-wounding (n= 3-4 mice per condition). Graphs represent averages of relative mRNA ± SEM. P values obtained Student’s T test.

### Supplemental Figure S3: Oscillatory expression of antiviral genes in mouse skin

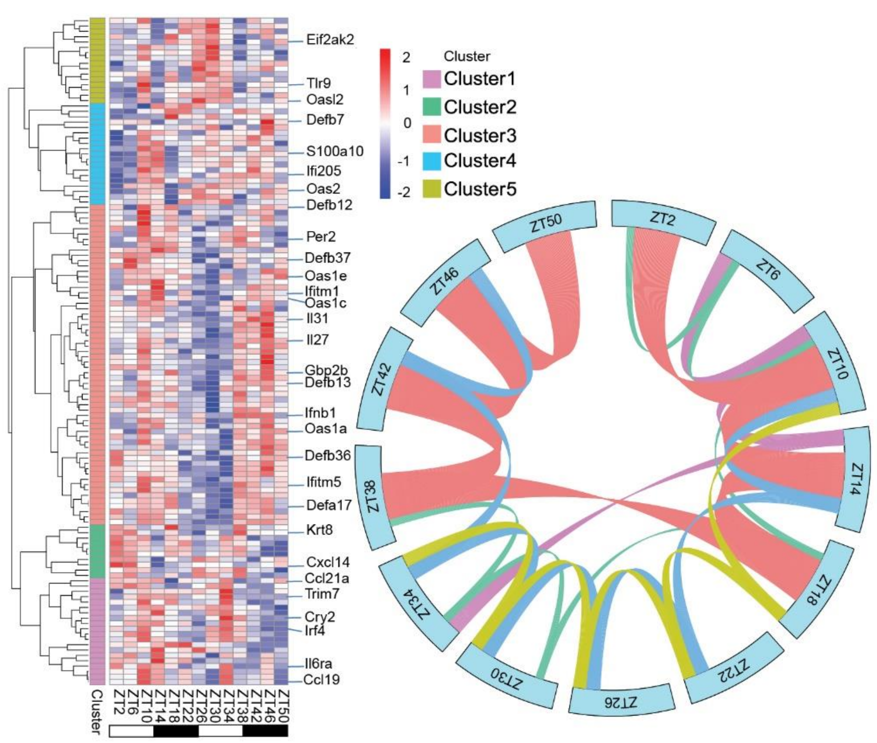


**Supplementary Figure S3:** CIRCOS plot displays oscillatory antiviral immune suppression of murine skin. Immune related genes were clustered into 5 groups based on timed expression data. Ribbon size indicates number of genes expressed at each time point.

**Supplemental Figure S4: Gating strategy for Figure 3c**

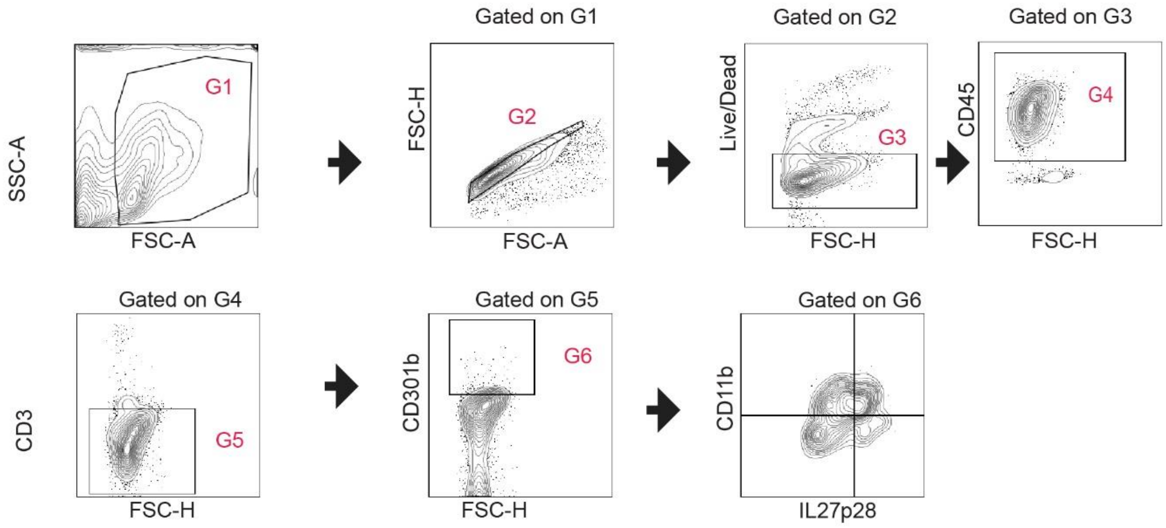


**Supplementary Figure S4:** Gating strategy for *Bmal1-/-* and wild type skin wounds. Circadian mutant and wild type skin were wounded, and flow cytometry was used to gate IL27 producing cells. Cells were pre-gated on live single cells that were CD45+, CD3+, CD301b+, and CD11b+.

**Supplemental Figure S5: Rhythmic AVP expression in primary human keratinocytes**

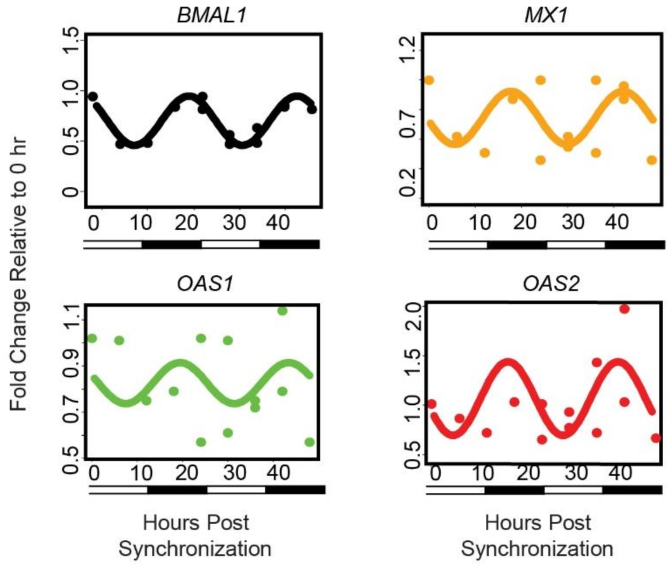


**Supplementary Figure S5:** Cosinor expression modelling of BMAL1, MX1, OAS1, OAS2 in human keratinocytes. Data from Figure 4A used to map and model a cosinor sine wave.

**Supplemental Figure S6: Circadian agent SR8278's antiviral effect is dependent on the presence of antiviral proteins in the skin**

a

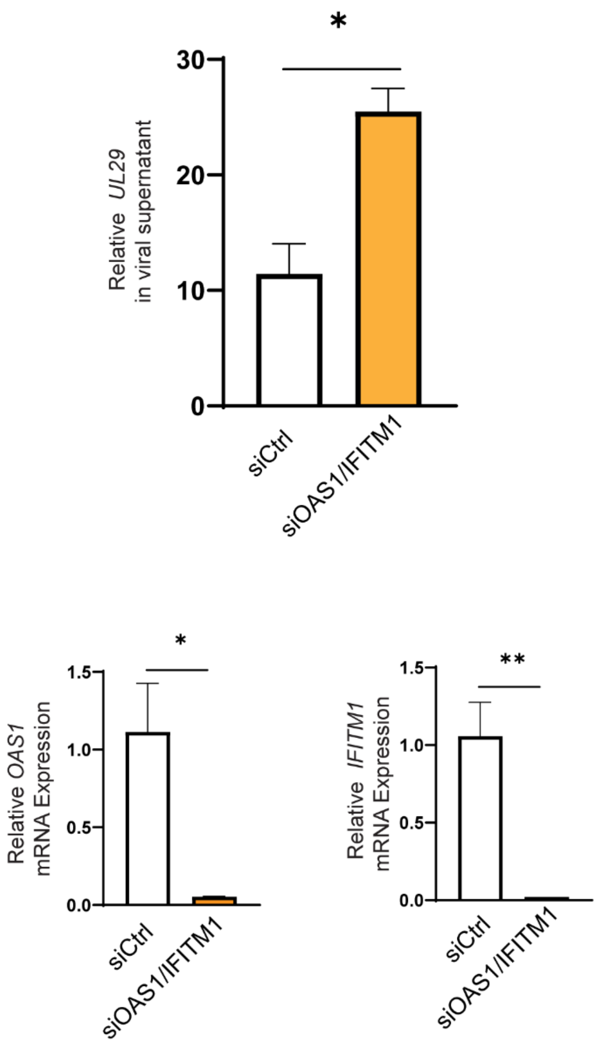


**Supplementary Figure S6:** NTERT keratinocytes were transfected with siRNA against siControl (Ctrl) or a combination of OAS1/IFITM1. Cells were infected with HSV1 at an MOI of 0.01 24 hours later, and treated with SR8278 at 10uM. Viral supernatant measurements were used against a standard curve of HSV1 UL29, and knockdown confirmed using RT-PCR. n=3 technical triplicates, representative of three experiments.

**Supplemental Figure S7: Circadian agent Nobiletin decreases HSV1 infection in human skin**

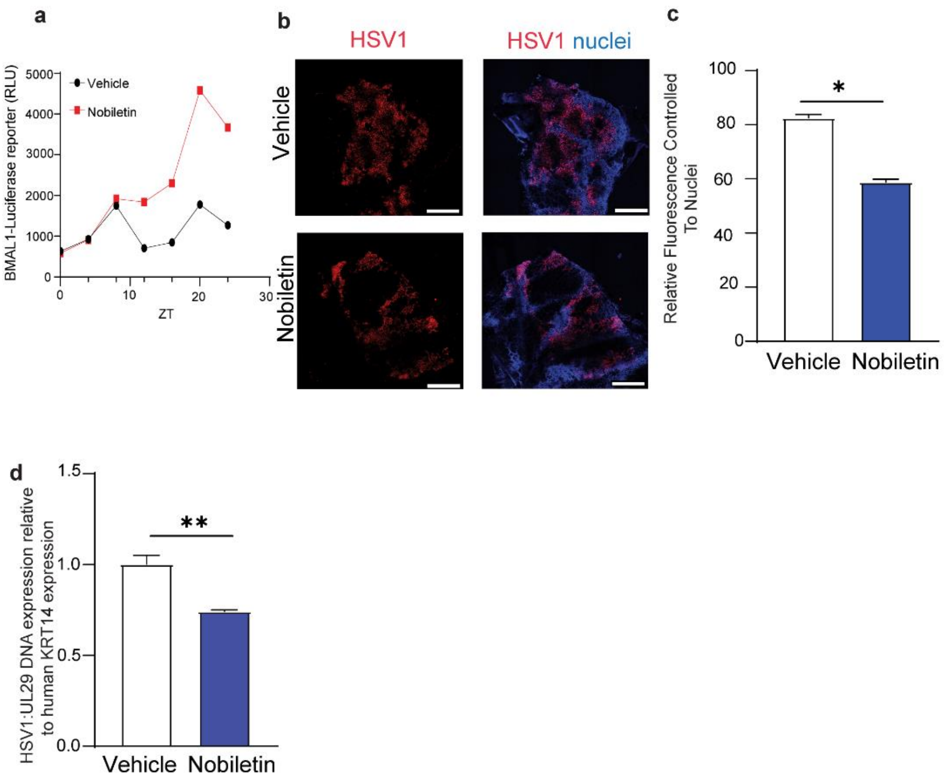


**Supplementary Figure S7:** Nobiletin can potentiate cutaneous circadian rhythms and reduce viral replication in the skin**. a)** Luciferase reporter assay. Keratinocytes transduced to express *Bmal*-luciferase were synchronized via overnight incubation with growth factor supplement omitted media and treated with vehicle or Nobiletin (5uM). Cells were harvested every 4 hours before luminescence measurements. **b)** Immunofluorescence of HSV in human skin treated with either vehicle or nobiletin (10uM). Bar = 500um. **c)** ImageJ (Fiji) quantification of relative viral immunofluorescence as controlled relative to nuclear staining. **d)** qPCR of HSV1 viral gene UL29 relative to human KRT14 in epidermal skin infection. Graphs represent averages of relative DNA ± SEM. P values obtained using Student’s T test.
